# Maternal behavior and the neonatal HPA axis in the Wistar Audiogenic Rat (WAR) strain: early life implications for a genetic animal model in epilepsy

**DOI:** 10.1101/2021.01.11.426211

**Authors:** Lívea Dornela Godoy, Norberto Garcia-Cairasco

## Abstract

Epileptogenesis is a multistage process and seizure susceptibility can be influenced by stress early in life. *Wistar Audiogenic Rat* (WAR) strain is an interesting model to study the association between stress and epilepsy, since it is naturally susceptible to seizures and present changes in the hypothalamus-pituitary-adrenal (HPA) axis activity. All these features are related to the pathogenic mechanisms usually associated to psychiatric comorbidities present in epilepsy. Therefore, the current study aimed to evaluate the neonate HPA axis function and maternal care under control and stress conditions in the WAR strain. Maternal behavior and neonate HPA axis were evaluated in Wistar and WAR strains under rest and after the presence of stressors. We observed that WAR pups present higher plasmatic corticosterone concentration as compared to Wistar pups. Although WAR dams do not show significant altered maternal behavior at rest, there is a higher latency to recover the litter in the pup retrieval test, while some did not recover all the litter. WAR dams presented similar behaviors to Wistar dams to a female intruder and maternal care with the pups in the maternal defense test. Taken together, these findings indicate that the WAR strain could show HPA axis disruption early in life and dams present altered maternal behavior under stressful events. Those alterations make the WAR strain an interesting model to evaluate vulnerability to epilepsy and its associated neuropsychiatric comorbidities.

## 1. INTRODUCTION

In many mammal species, maternal care requires the recruitment of a wide range of behavioral and physiological adaptations. Postpartum rodents are highly motivated to care for, nourish and defend pups [1]. Several experimental models that alter maternal care early in life have demonstrated how this disruption promotes stress and neuroplasticity that are related to psychopathologies [2–4].

Further evidence also points out that stress is relevant to the process of epileptogenesis, both in childhood and in adult life. Epileptogenesis is a multi-stage process, which can begin early in life and may be influenced by stress [5]. It is currently suggested that early stress may create a permanent vulnerability to the development of epilepsy [6]. The effects of stress on the brain are mediated by several types of molecules, including neurotransmitters, peptides and steroid hormones, which are candidates for mediating the influence of early life stress on general network excitability [6–8]. Only a few reports dedicated to the investigation on the impact of early life stress and were able to make the link with stress-induced epileptiform discharges, mainly in limbic regions, suggesting its possible role on the seizure propagation and contributory role on epileptogenesis [7,9–15]. None of those studies has assessed the relationship between postnatal early life stress and epilepsy in seizure prone strains, with the exception of the audiogenic and absence seizure models in WAG/Rij and absence seizure in GAERs strains [16]. Glucocorticoids play a contributory role in the epileptogenic process [6,17,18], through several actions in the brain that could mediate this outcome, such as changes in the networks associated with seizures, or their direct effects on the excitability of the limbic system [5]. Therefore, hyperactivity in the hypothalamus-pituitary-adrenal axis (HPA), resulting from either stress or exogenously applied corticosterone, primes the brain for epilepsy [19].

Most experimental models study stress outcomes in pups by either limiting the resources or artificially reducing maternal care [20]. Only a few studies were able to relate the disruption early in life with epilepsy in a naturalistic way (not an artificially imposed intervention like maternal deprivation/separation, but through protocols involving a species related context and a translational perspective) [20,21] or by monitoring previous maternal behavior dysregulation in models with specific genetic background [7,16,22,23].

In this context, we aim to study the Wistar Audiogenic Rat (WAR), an inbred strain that has been genetically selected through the mating of Wistar (sisters x brothers) rats that present behavioral seizures when exposed to acoustic stimuli [24–26]. WAR is considered as a genetically selected reflex model, since in face of the acoustic stimulus animals present brainstem-dependent tonic–clonic seizures, and with the acoustic stimuli repetition (audiogenic kindling) is capable of propagate and recruit limbic regions [27]. In addition, during adult life, WARs show HPA axis disruption represented by hyperplasic adrenal gland, disruption of the circadian cycle for adrenocorticotrophic hormone (ACTH) secretion and higher plasma corticosterone levels in response to exogenous ACTH [28]. Moreover, animals from this strain present higher levels of anxiety [29] and depressive like-behaviors [30]. Therefore, considering that an extensive body of evidence suggests the important role of maternal care in the proper development of the central nervous system, as well as changes in maternal care associated to neurological and psychiatric disorders, we propose that the WAR strain may be an interesting model for the study of maternal care and the early life stress role in epileptogenesis and neuropsychiatric comorbidities associated with epilepsy.

## 2. METHODS

### 2.1 Animals

Wistar and WARs were acquired from the Central Animal Facility and from the Special Rat Strains Facility, respectively, both from the University of São Paulo, Ribeirão Preto Campus. Animals were kept in transparent acrylic boxes (45 x 32 x 17 cm) and housed in the maintenance room of the Department of Physiology, under controlled ventilation and temperature (25 ± 2 °C), with a light/dark cycle of 12 hours (lights at 7:00 AM). Animals were allowed free access to water and food. This study was approved by the Ethics Committee on Animal Experimentation (CETEA) of the Medical School of Ribeirão Preto - USP (80/2014). Different litters were assigned to either hormonal or behavioral experiments.

#### 2.1.1. Mating, Gestation and Birth

At 75 days of age, one male for every two nulliparous females was mated (Wistar n=14 and WAR n=13). Pregnancy was confirmed by checking the week following mating for: (i) vaginal plug (which is the seminal fluid from the male that prevents the female from mating with another male), (ii) significant body mass increase and (iii) changes in nipples. Pregnant females of each strain were kept in individual cages in conditions of controlled temperature and luminosity and maintained as previously mentioned. On the day of birth (Post natal day 0 - PD0) litters were ideally culled on eight pups with equally distributed sexes. Birth parameters such as the number of pups, sex distribution and possible malformations or miscarriages were recorded.

### 2.2 Plasmatic Corticosterone at rest and after stress

At PD12, Wistar and WAR offspring (from 2 different litters for each group) were decapitated under basal condition or after a physical stressor (n=6-12). The physical stressor protocol was performed according to Godoy et al [31], in which pups received a subcutaneous (s.c.) injection of propyleneglycol volume of 0.1 ml. After injection animals were returned to their home cages. After 30 min the animals were decapitated and trunk blood was collected (between 8:00 AM and 9:00 AM) for plasmatic corticosterone assessment.

Briefly, plasma was separated by centrifugation at 4°C and was stored in a freezer at −20°C. The hormone concentration was determined by a radioimmunoassay protocol [28,31,32].

### 2.3 Maternal behavior in Wistar and WAR strain

#### 2.3.1 Maternal care

Maternal behavior (Wistar n=8; WAR n=9) was monitored for 60 min each, from PD1-PD9. Records occurred at regular periods, daily in the light phase period (08:00 AM - 10:00 AM). In each observation period, dam behavior (Supplementary Table 1) was recorded every 2 min (30 observations per session). Scores were calculated by dividing the behavior frequency by the total number of observations. Also, the behaviors were considered if occurred *in* or *off* the nest.

#### 2.3.2. Pup retrieval test

In this test, at PD2, dams were removed from their home cage and were placed in another room for 30 min. Meanwhile, pups were distributed in the home cage, away from the nest [1,33]. After this period, the dam was returned to the home cage, and behaviors were recorded on video for 30 min. Two blind experimenters assessed the latency for retrieving the first pup, half litter and the whole litter. It is important to mention that the litter size presented a small variation (mean number of pups/litter = 7.2; Standard Deviation = 1.135).

#### 2.3.3. Maternal defense test

The maternal defense test was performed at PD6. Animals were transported to the experimentation room 60 min before the test. Dams were confronted with a virgin and unknown intruder (from the same strain) in the home cage at the presence of the litter for 10 min [1,34]. An experienced observer, blind to the experimental group, analyzed the video recordings. Behavioral parameters are described in Supplementary Table 2.

### 2.4. Statistical analysis

All analyses were performed with GraphPad Prism 7.0 (EUA). Hormonal data were analyzed with 2-way ANOVA (strain and stress factors). Birth and behavioral data were analyzed with Student’s t-test (two-tailed) for comparison between the two groups. Maternal behaviors and time in nest from PD1-PD9 were also analyzed by 2-way repeated measures ANOVA (strain and time factors). Additionally, in the pup retrieval test, Chi-square analysis of dams that fully retrieved their litter and matrix correlation (Pearson correlation coefficients between the possible pairs of variables) of dams litter size, latencies to retrieve first pup, half litter and all litter were compared. It was considered p <0.05 as a significant difference.

## 3. RESULTS

### 3.1 Mating, Gestation and Birth

In 14 Wistar matings, 10 females gave birth, and, from those, there was one dam that presented cannibalism and was removed from the study. In 13 WAR matings, 11 females gave birth. There was cannibalism in one litter and, another dam gave birth to two pups, so both were removed from the study. Comparing the birth rate between both strains, we observed no significant difference in the number of pups at birth (t=1.61 df=15; p=0.12) (Figure 1).

**Figure 1.**
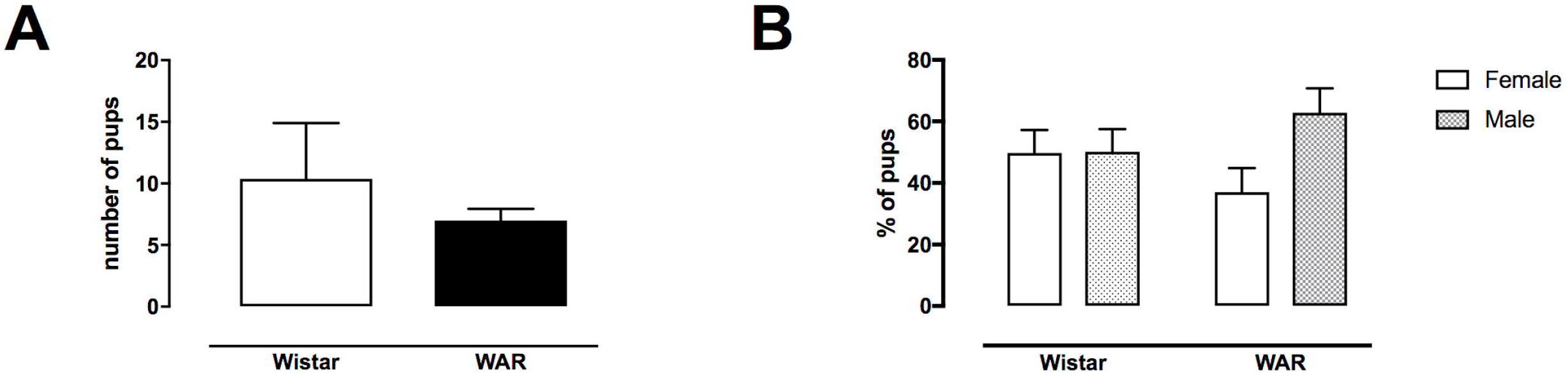
Birth parameters from Wistar and WAR nulliparous dams (n=8-9). A. Number of pups at birth (Mean ± SEM 10.38 ± 1.6 and 7 ± 0.9, respectively). B. Offspring sex distribution. (Mean ± SEM for Wistar males 50.2% ± 7.3 and females 49.8% ± 7.4; Mean ± SEM for WAR males 63% ± 7.8 and females 37% ± 6.8).

### 3.2 Plasmatic Corticosterone at rest and after stress

In the corticosterone assay, there is a significant difference for strain (F[1,26]= 9.09; p=0.008), stress (F[1,26]= 8.234; p=0.005) but not for interaction (F[1,26]= 0.75; p=0.39) (Figure 2).

**Figure 2.**
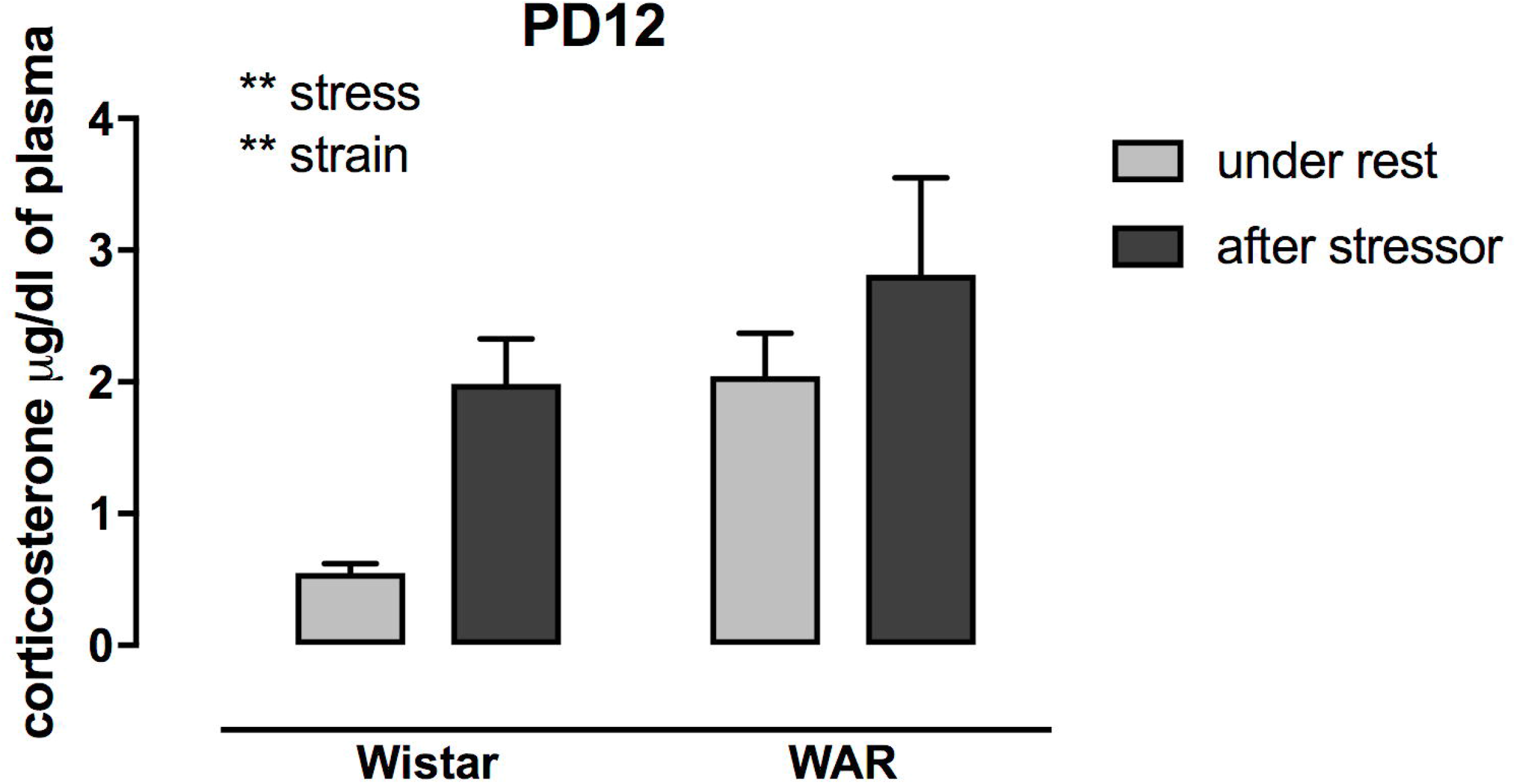
Plasmatic corticosterone concentration in Wistar and WAR rats under rest (grey) or 30 min after a physical stressor (black) (n=4-12 per group). There is a significant difference for strain and for stress (**p<0.01), but not for interaction. Under rest (Mean ± SEM) for Wistar (0.55 ± 0.06) and WAR (1.98 ± 0.34). After stressor (Mean ± SEM) for Wistar (2.04 ± 0.32) and WAR (2.81 ± 0.73).

### 3.3 Maternal care

All maternal and non-maternal behaviors were represented in the chart, demonstrating there is no overall difference between WAR and Wistar behavior, collapsing analysis from PD1-PD9 (Figure 3 and Supplementary Table 1).

**Figure 3.**
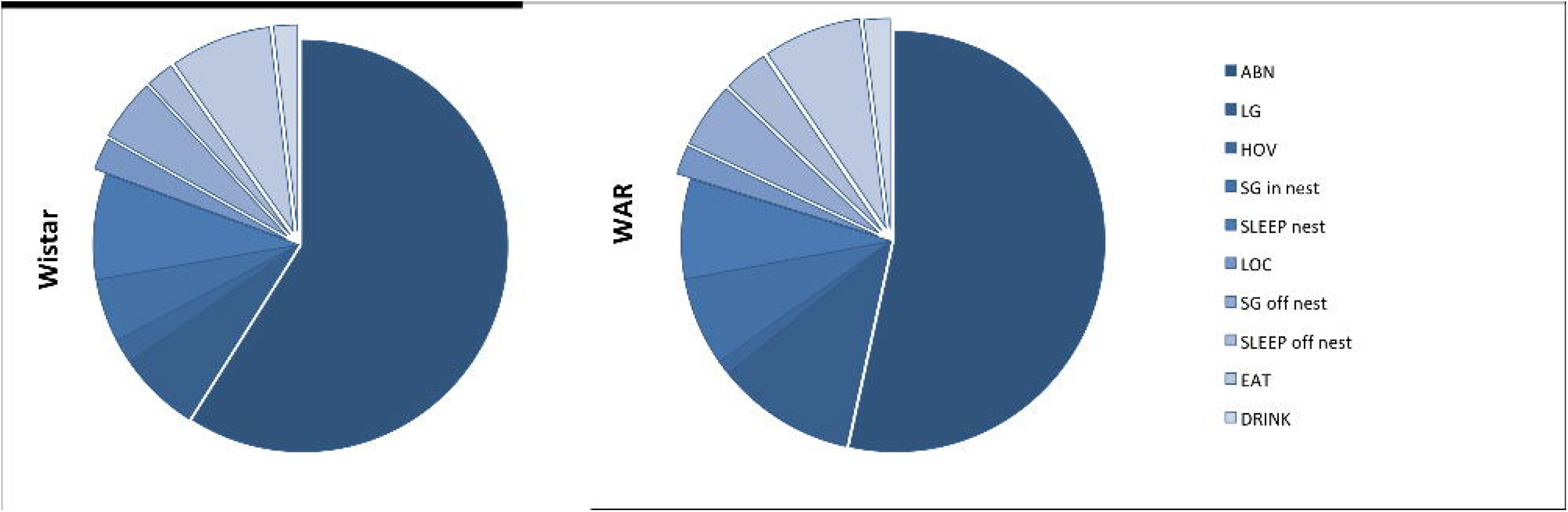
Representative graph of the mean of all behaviors in Wistar (left) and WAR strain (right) from PD1 to PD9 (n=8-9). Dark blue behaviors occurred in the nest and taller light blue off the nest. ABN-Arched back nursing; LG-Licking Grooming; HOV-Hovering; SG-Self grooming; SLEEP-sleeping; LOC-locomotion; EAT-eating; DRINK-drinking.

In Figure 4, the main maternal behaviors at PD1 and from PD1 to PD9 are represented. At PD1 there were no differences in the number of arched back nursing posture (ABN; t=0.04; df=15; p=0,96) and time spent in the nest (t=0.63; df=15; p=0.53) (Figure 4A and 4C). However, in PD1, WAR dams exhibited higher licking and grooming (LG) when compared to Wistar (t =2.62; df=15; p =0.01; Figure 4B).

**Figure 4.**
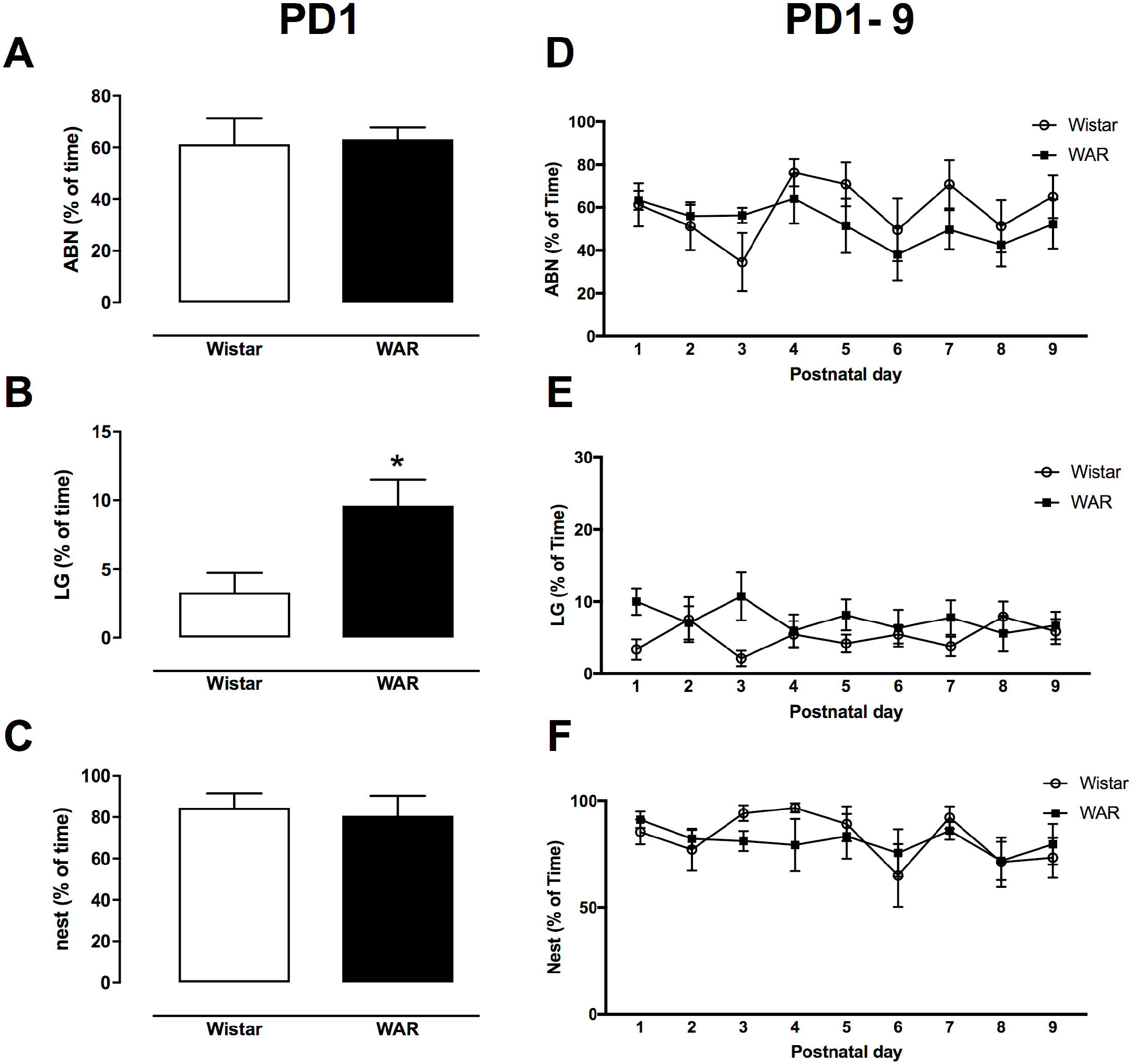
Maternal behaviors in Wistar (white) and WAR (black) strains under undisturbed conditions in their home cage on PD1 (left column) and during PD1-to PD9 period (right). A,D. Arched Back Nursing (ABN) posture. B,E. Licking and Grooming (LG). C,F. Dam in the nest. At PD1 (Mean ± SEM) for Wistar and WAR, respectively in ABN (57 ± 5 and 61 ± 5); LG (3 ± 1.4 and 9 ± 1.8); Dam in nest (84 ± 6 and 80 ± 9); *p<0.05 compared to Wistar at PD1. From PD1-to PD9 period (right) there is no significant difference for strain, time, nor for interaction (n=8-9).

As demonstrated previously in the graph chart, during the PD1 to PD9 period (Figure 4 D-F) there was no strain effect for ABN (F_(1,15)_=1.01; p=0.32), LG (F_(1,15)_=3.63; p=0.07) or total time in the nest (F_(1,15)_=0.095; p=0.76). There were no significant differences for time in ABN (F_(8,120)_=1.62; p=0.12), LG (F_(8,120)_=0.12; p=0.99) and for total time in the nest (F_(8,120)_=1.72; p=0.1) or interaction in ABN (F_(16,120)_=0.93; p=0.48), LG (F_(16,128)_=1.48; p=0.17) and total time in the nest (F_(16,120)_= 0.69; p=0.69).

### 3.4 Pup retrieval test

In the pup retrieval test (Figure 5), WAR presented higher latencies to group the first pup (t = 2.159 df = 8; p = 0.06), reaching significance at half of the litter (Figure 5B) (t = 2,625 df = 7, p = 0.03) and whole litter retrieval (Figure 5) (t = 2.473 df = 7, p = 0.04). For instance, 80% of Wistar dams grouped the whole litter, whereas in WAR strain they were 55% (χ^2^= 8.36, df = 1, p = 0.36). Matrix correlation showed that there are no significant differences for litter size and latency to first pup (r=-0.32; p=0.39), latency to half litter (r=0.10; p=0.7811) or latency to all litter (r=0.21; p=0.58).

**Figure 5.**
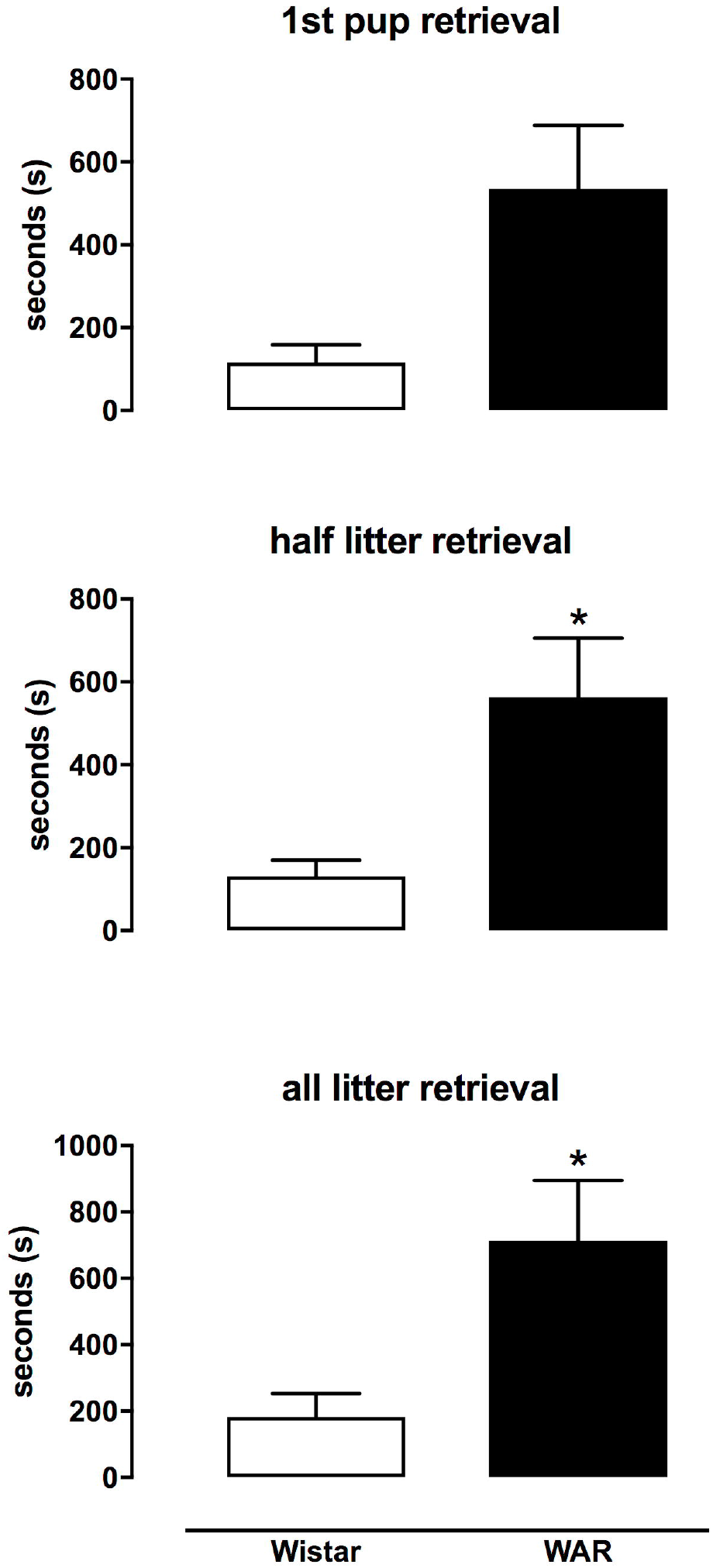
Behavioural assessment of Wistar (white) and WAR (black) dams in pup retrieval test. A. Latency to retrieve the first pup (Mean ± SEM) for Wistar and WAR, respectively (116 ± 42 and 535 ± 152). B. Latency to retrieve half litter (Mean ± SEM) for Wistar and WAR, respectively (131 ±39 and 564 ± 142). C. Latency to group all the litter (Mean ± SEM) for Wistar and WAR, respectively (182 ± 70 and 713 ± 181). *p<0.05 compared to Wistar.

### 3.5 Maternal Defense test

In the maternal defense test (Supplementary Figure 1), there was no significant difference in number of attacks (t = 0.87; df = 11; p = 0.66), latency for the first attack (t = 0.73; df = 10; p = 0.47), aggressive behaviors (t = 0.05; df = 11; p = 0.95) or threat behaviors (t = 0.02; df = 11; p = 0.98). The latency and frequency of ABN posture were also evaluated after the maternal defense test (Supplementary Figure 2). It was observed no significant difference in the frequency (t = 1.70; df = 11; p = 0.84) and latency (t = 0.38; df = 11; p = 0.70) when comparing the two strains.

## 4. DISCUSSION

In the current study we demonstrated that the dams of the WAR strain have no difference in the number of pups born, when compared to dams of the Wistar strain and there was no evidence of any malformation. Previous data from our laboratory confirm these findings [35]. Although WAR animals do not differ in number of born pups as compared to Wistar [35], a study from our group revealed that the WAR strain presented hyperproteinemia, hypertriglyceridemia, and embryonic losses suggesting an inadequate intrauterine condition for embryonic development and fetal viability [36]. WARs present both reduced weight at birth and reduced body weight gain during the lactation period and adulthood [28,35] and a recent study revealed a higher percentage of skeletal anomalies in fetuses compared with Wistar [36].

Compared to Wistar, animals of the WAR strain show a higher plasmatic concentration of corticosterone, irrespective of the presence of a stressor under rest condition. Briefly, data from our laboratory have already demonstrated that, compared to Wistar, male adult WARs show many altered features in HPA axis later in life, such as increased corticosterone during the evening period, increased response to restraint stress and to ACTH stimulation; increased adrenal mass (higher fasciculata cortical layer area); and lower body mass [28]. WARs display increased anxiety- and depressive-like behaviors [30,37], and cognitive deficits [30] - behavior alterations that also have been implicated as comorbidities associated to stress system dysfunction [38], which contributes to the important discussion on epilepsy comorbidities [19,39]. The WAR strain was derived from the inbred selection of Wistar rats that basally displayed seizure susceptibility to acoustic stimulation. The various strain differences were evaluated by many research laboratories, in collaborative studies, involving other audiogenic seizure prone strains and also other models of epilepsy [27]. Studies involving the WAR strain and many of the alterations were discussed and further reviewed [27]. Considering this scenario, more components of the HPA axis must be evaluated to understand how the maternal behavior, offspring corticosterone expression compose this complex and intricate stress system in this audiogenic seizure-prone strain. Thus, our present data add to the discussion of HPA axis changes in a life-span perspective of the WAR strain.

Specific maternal behaviors during the first postnatal weeks have been associated with changes in neurodevelopment, especially HPA axis programming [20,40]. Manipulations that alter those components are responsible for 1) an inhibitory effect that maintains low corticosterone levels and 2) a suppressing effect that prevents the corticosterone response from being activated under stress, once the pup is capable of responding [41]. This feature is well known as Stress Hyporesponsive Period (SHRP) and is maintained by specific components of maternal care, such as LG and milk delivery, and maternal behavior disruption and stressful situations may contribute to disturbances in this sensitive period [42–45].

Even though there are no major differences in maternal behavior under rest conditions, we observed changes in the pup retrieval test, since WAR dams failed to group the litter, demonstrated by higher latencies to retrieve pups and not grouping the whole litter. Maternal care in the postpartum period has an important component represented by direct behaviors towards the pups associated with litter demands [46–48]. In the WAG/Rij strain, an absence and audiogenic seizure prone rat strain, higher latencies to retrieve pups and fewer approaches are also detected [23]. In another strain susceptible to epilepsy, the EL mouse, changes in maternal behavior were also reported. It has been shown that in this strain, the EL females exhibited an increased latency in nestling the pups, which may also reflect a poorly adaptive response to a new environment, as well as a change in the motivational system [49].

Behavioral changes that emerge after the postpartum period are mediated by a wide variety of neurotransmitters such as dopamine and neurohormones such as oxytocin (OT) and ACTH. WARs show higher ACTH secretion [28] and previous studies from our laboratory have suggested that WARs, could present alterations in the OT system [50], which might be related to a naturally excessive grooming when exposed to a novel environment. As a consequence, it would be interesting to study neurohormonal brain circuits in WARs and verify how they contribute to specific behaviors directed to the pup care, like LG and retrieving.

Adding to the WAR maternal behavioral monitoring, we did not observe significant changes in the maternal defense test. It is important to consider that, although maternal care represents a strong relationship between the mother and the offspring, maternal aggression is an activity directed toward an adult intruder [55]. The nature of the two behaviors is different and consequently involves different neural circuits.

Thus, the importance of approaching a multifactorial view is required on studying epilepsy models [49], such as the current one with the WAR strain. Parental investment and early life stress can affect seizures and behavior in adult offspring phenotype [49]. Therefore, perspectives that include both neurodevelopment and environmental studies in genetic models, as opposed to exclusively investigating models with fully and regular developed adult animals, may provide further advances to make inferences about epileptogenesis, which is a phenomenon extremely affected by early life experiences [52,53].

## 5. LIMITATIONS

In this study, the corticosterone levels were assessed using two litters per group at only one time point. It would be interesting in future studies to include more time points, not only confirming this disruption in HPA axis but also evaluating corticosterone considering the circadian rhythm.

## 6. CONCLUSION

The current data indicate that the WAR strain does not show differences under rest conditions, but present maternal behaviors alterations in the face of stressful events. It is possible that the altered maternal behaviors in the WAR strain are specifically related to those directed towards the pups and are not involved with defense/aggression behavior. In addition, the WAR strain shows an increased plasmatic corticosterone concentration early in life, adding to previous data on increased HPA axis activity, which may contribute to the epileptogenic process and to stress-related psychiatric comorbidities already described in this genetically selected strain.

## 7. ACKNOWLEDGMENTS

The authors would like to thank Jose Antonio Cortes de Oliveira for the assistance with the breeding and maintenance of the WAR colony and the valuable discussions, and Palloma Beatriz Fialho Damas for helping with double blind analysis.

## Abbreviations

ABN: arched back nursing posture
ACTH: adrenocorticotrophic hormone
CORT: corticosterone
HPA: hypothalamus-pituitary-adrenal axis
LG: Licking and Grooming
OT: Oxytocin
PND: Post Natal day
WAR: Wistar audiogenic rat.

---

**Supplementary Table 1.** Parameters considered on behavioural assessment of Maternal Care, adapted from [54]. The means % of time of all behaviors from P1-P9 represented in Figure 3 are detailed

| **Observation of Maternal care (P1-P9)** | **Wistar (Mean%)** | **WAR (Mean%)** |
| --- | --- | --- |
| arched back nursing posture (ABN) | 62 | 57 |
| licking grooming (LG) | 5 | 8 |
| Hovering (HOV) | 2 | 1 |
| self grooming (SG) in nest | 5 | 7 |
| sleeping (SLEEP) in nest | 8 | 8 |
| locomotion (LOC) | 3 | 2 |
| self grooming (SG) off nest | 5 | 5 |
| sleeping (SLEEP) off nest | 2 | 3 |
| Eating (EAT) | 8 | 7 |
| Drinking (DRINK) | 2 | 2 |

**Supplementary Table 2.** Parameters considered on Pup Retrieval and Maternal Defense tests.Figure Captions

| **Pup Retrieval Test** | **Maternal Defense Test** |
| --- | --- |
| Latency for recovery of the first pup | total number of attacks |
| Latency for recovery of half litter | latency for the first attack |
| Latency for recovery of half litter | Aggressive behaviors that include contact with the intruder or biting |
|  | Lateral and vertical threat |
|  | Latency and % ABN after the end of test |

**Supplementary Figure 1.**
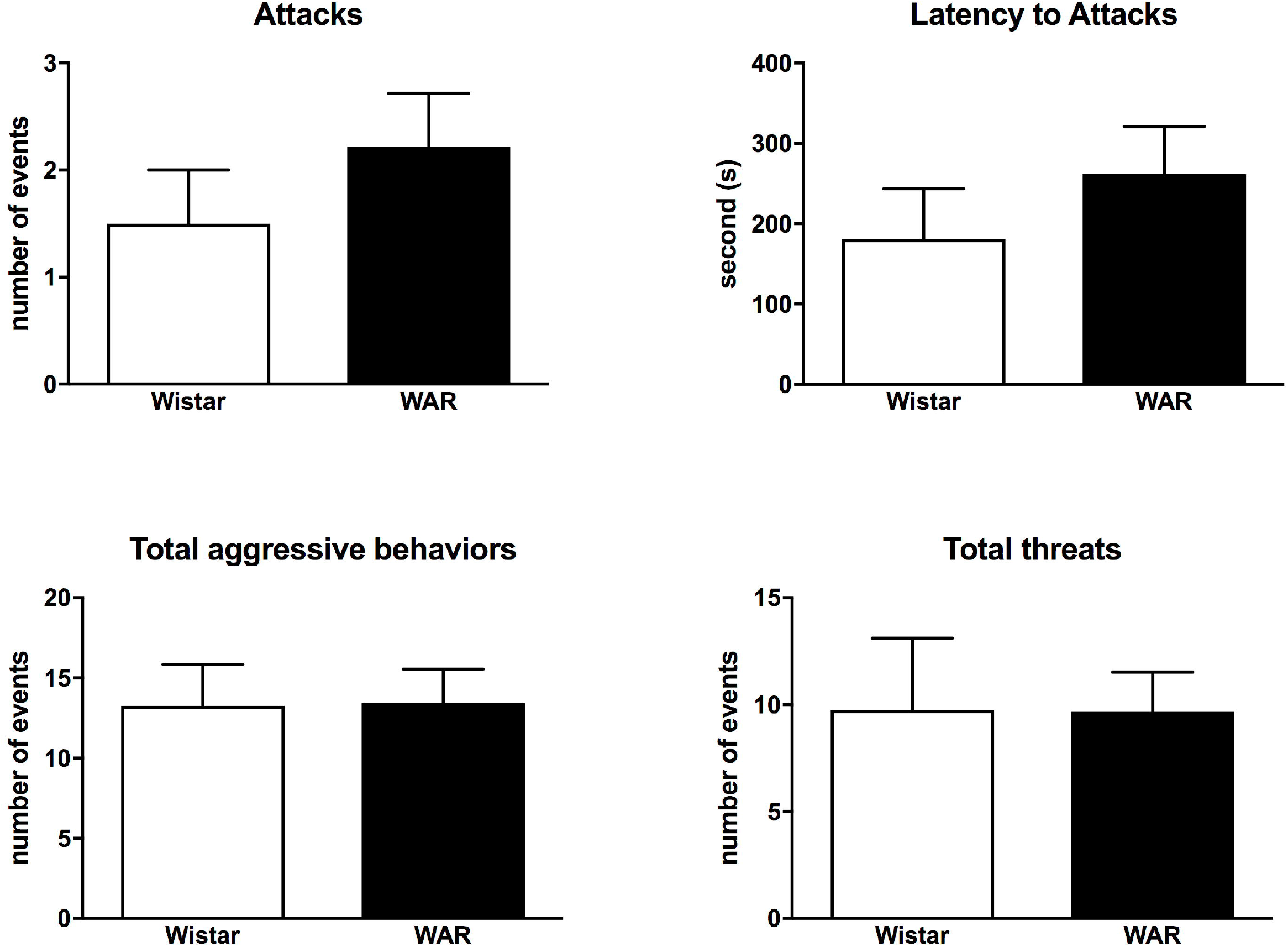
Evaluation of Wistar (white) and WAR (black) dams behaviors in maternal defense test, upon exposure to unknown virgin untruder in the homecage (n=8-9). A. Attack Behavior (Mean ± SEM) for Wistar and WAR, respectively (1.5 ± 0.5 and 2.2 ± 0.5) B. Latency for first attack (Mean ± SEM) for Wistar and WAR, respectively (180 ± 62 and 262 ± 59) C. Aggressive behavior (Mean ± SEM) for Wistar and WAR, respectively (13 ± 2.5 and 13 ± 2) D. Threat Behavior (Mean ± SEM) for Wistar and WAR, respectively (9 ± 3 and 9 ± 1.8).

**Supplementary Figure 2.**
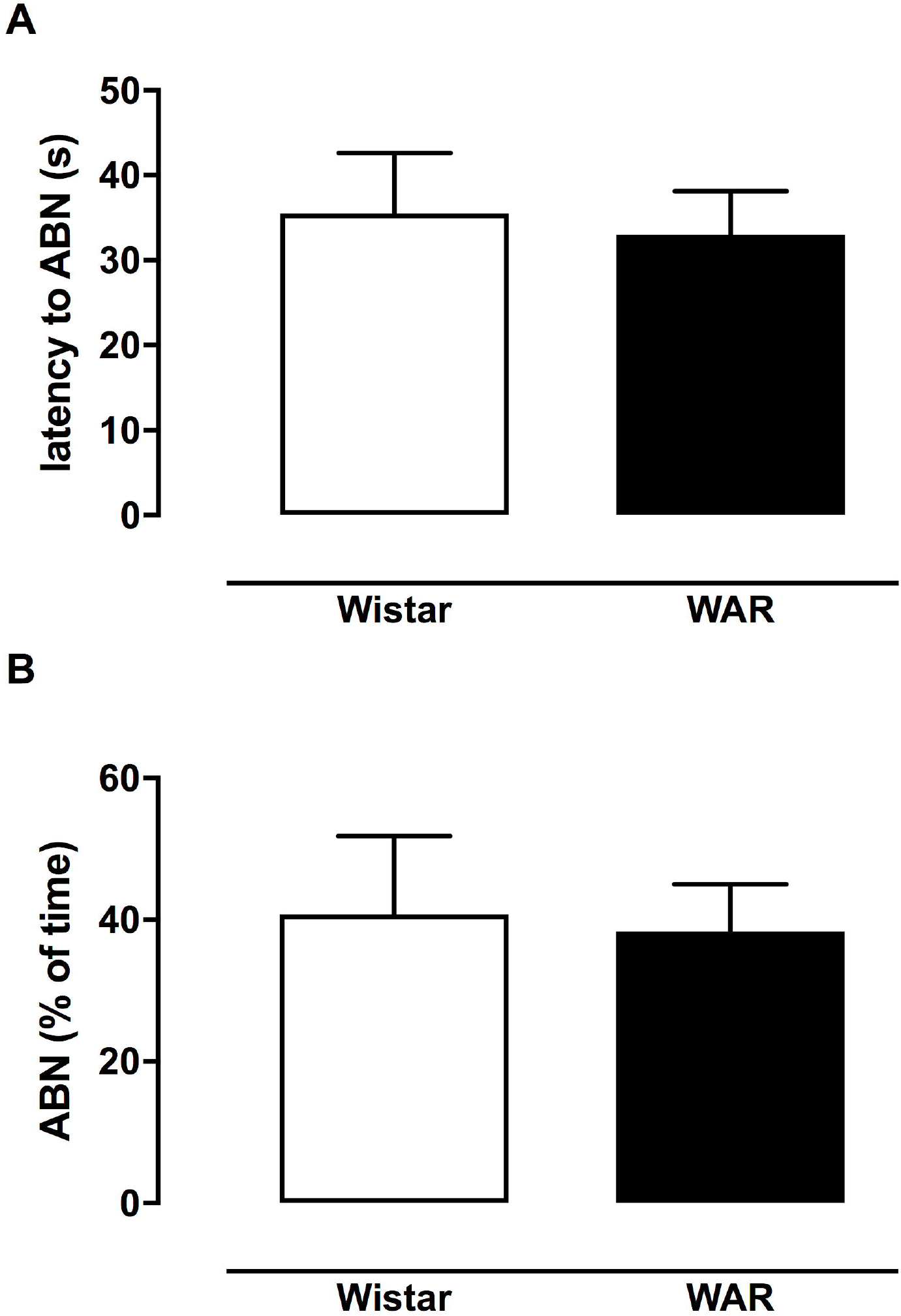
Evaluation of Wistar (white) and WAR (black) Arched Back Nursing (ABN) after maternal defense test (n=8). A. Latency (Mean ± SEM) for Wistar and WAR, respectively (35 ± 7 and 33 ± 5) and B. Frequency (Mean ± SEM) for Wistar and WAR, respectively (40 ± 11 and 38 ± 6).

